# Serotonin drives aggression and social behaviours of laboratory mice in a semi-natural environment

**DOI:** 10.1101/2024.02.02.578690

**Authors:** Marion Rivalan, Lucille Alonso, Valentina Mosienko, Patrik Bey, Alexia Hyde, Michael Bader, York Winter, Natalia Alenina

**Author notes:** Corresponding authors and present e-mail addresses: Marion Rivalan E-mail address, Natalia Alenina. Authors contributed equally to this work (York Winter, Natalia Alenina).

## Abstract

Aggression is an adaptive social behaviour crucial for the stability and prosperity of social groups. When uncontrolled, aggression leads to pathological violence that disrupts group structure and individual well-being. The comorbidity of uncontrolled aggression across different psychopathologies makes it a potential endophenotype of mental disorders with the same neurobiological substrates. Serotonin plays a critical role in the regulation of impulsive and aggressive behaviours, and mice lacking brain serotonin, due to the ablation of a rate-limiting enzyme of serotonin synthesis (Tryptophan hydroxylase 2, TPH2), are a potential model of pathological aggression. Home cage monitoring allows for the continuous observation and quantification of social and non-social behaviours in group-housed, freely-moving mice. Using an ethological approach, we investigated the impact of central serotonin ablation on everyday expression of social and non-social behaviours and their correlations in undisturbed, group-living *Tph2*-deficient and wildtype mice. By training a machine learning algorithm on behavioural time series, “allogrooming”, “struggling at feeder” and “eating” emerged as key behaviours dissociating one genotype from the other. Although *Tph2*-deficient mice showed characteristics of pathological aggression and decreased communication compared to wildtype animals they still showed affiliative behaviours to normal levels. Altogether, such distinct and dynamic phenotype of Tph2-deficient mice influenced the group’s structure and the dynamic of its hierarchical organization which emerged later. These aspects were analyzed using social network analysis and the Glicko rating methods. This study demonstrates the importance of the ethological approach for understanding the global impact of pathological aggression on different aspects of life both at the individual and the group level. Home cage monitoring allows the observation of the natural behaviours of the mice in a semi-natural habitat and provides an accurate representation of real-world phenomena and pathological mechanisms. The results of this study provide insights into the neurobiological substrate of pathological aggression and their potential role in complex brain disorders.

## Introduction

Aggression is an adaptive behaviour, often the result of competition (Nelson and Trainor 2007). It is an important social behaviour that ensures the stability and the prosperity of a social group (Van Loo et al., 2003, Sapolsky et al., 2005, Wang et al., 2011). When it cannot be avoided, aggression is typically short and directed to acquire and keep resources including territory, mating partners, and food (Kiser et al., 2012). On the pathological side escalated aggression or violence, excessively and repeatedly hurting others or the perpetrator itself, happens in all contexts and is devoid of a communication purpose (Natarajan et al., 2009). This extreme behaviour is detrimental to the individual as it can result in death or invalidity for example, and it severely disrupts the group’s structure, its security and comfort (WHO 2004). Interpersonal violence has a high economical cost and a better understanding of how aggression impacts group structure is critical for “diagnosis, prevention, and treatment, but also for guidance of public and judicial policies” (WHO 2004, Miczek et al., 2007).

Uncontrolled aggression and violence are diagnostic criteria of different psychiatric disorders, such as schizophrenia, alcoholism, intermittent explosive disorder, autism or dementia (Lesch et al., 2012, Volavka et al., 2008). Considering a dimensional and trans-diagnostic view of mental disorders, the comorbidity of uncontrolled aggression across different psychopathologies makes it a good potential endophenotype of mental disorders (Niederkofler et al., 2016, Nestler and Hyman 2010) with the neurobiological (and heritable) substrates of pathological aggression being the same across different psychiatric disorders (DSM-5, Gottesman et al., 2003, Gould et al., 2006, Robbins et al., 2012). Research into neurobiological substrates underlying aggression is thus essential to our insight into the etiology of complex brain disorders (Kalueff et al., 2015) and its potential global impact on society.

Serotonin (5-hydroxytryptamine, 5-HT) is a monoamine that plays a critical role in the regulation of impulsive and aggressive behaviours in humans and animals. Mice with a congenital lack of serotonin in the brain due to the lack of a rate-limiting serotonin-producing enzyme Tryptophan hydroxylase 2, TPH2 (*Tph2*-deficient mice, Alenina et al., 2009) were found to be very belligerent animals. They show higher levels of aggression toward strangers, poor social recognition abilities and an impulsive-like phenotype (Angoa-Pérez et al., 2012). *Tph2*-deficient mice also present behavioural abnormalities similar to human symptoms of Autistic Syndrome Disorder (ASD) and Impulsive related disorders (Mosienko et al., 2012, 2015a, 2015b, Kane et al., 2012, Beis et al., 2015, Angoa-Pérez et al., 2012, Kästner et al., 2019). In line with the search for trans-nosological symptoms of mental and neurological disorders, the *Tph2*-deficient mice thus represent a potential interesting model of pathological aggression.

The Visible Burrow System (VBS) is a semi-natural habitat first developed in rats (Blanchard and Blanchard, 1989) and more recently in mice (Arakawa et al., 2007, Pobbe et al., 2012, Bove et al., 2018) where ethological aspects (e.g. day/night fluctuation of activity spatial distribution, place preference) and social and non-social behaviours of group-housed, freely-moving individuals can be continuously observed and quantified (Alonso et al., 2020, 2023).

In an effort to model real world phenomena and pathological mechanisms reminiscent of everyday-life of human patients in mice (McCloskey et al., 2011), we chose to apprehend the individuals’ behaviours and the group dynamic of *Tph2*-deficient (*Tph2*^−/−^) and wildtype (*Tph2*^+/+^) mice directly in their housing environment. To this end, a new version of the VBS was designed and built. The aim of this study was to investigate the impact of central serotonin loss on everyday expression of social and non-social behaviours and their correlations over days in undisturbed, group-living mice of the same-genotype. Training a machine learning algorithm (Random Forest classifier, Breimann et al., 2001) on this extended behavioural data allowed us to identify key variables dissociating one genotype from the other when living in such ethologically-relevant environment. Because excessive aggression does not only drastically affect the life of the perpetrator but simultaneously affects the dynamic of the group and its structural organization, we evaluated if and how lack of central serotonin influenced the group’s structure and dynamic of its hierarchical organization using social network analysis (SNA, Krause et al., 2010) and the Glicko rating methods (Glickman et al., 1999), respectively.

## 2. Materials and Methods

### 2.1 Animals

Mice were maintained at the Max Delbrück Center (MDC) animal facility in individually ventilated cages (Tecniplast, Italy) under specific pathogen-free, standardized conditions in accordance with the German Animal Protection Law. Mice were group-housed at a constant temperature of 21 ± 2°C with a humidity of 65 ± 5%, and under 12h/12h light/dark cycle (light off at 18:00) and had *ad libitum* access to food and water throughout the project. Ten groups of four *Tph2*-deficient (*Tph2*^−/−^) or wildtype (*Tph2*^+/+^) male mice (*n_total_*=40) were used in this project: five groups of *Tph2*^+/+^ and five groups of *Tph2*^−/−^ mice on C57BL/6N genetic background (Alenina et al., 2009, Mosienko et al., 2012). An independent cohort of C57BL/6N male mice (*n*=6) was used as unfamiliar mice in the Three-chamber test.

### 2.2 Experimental design

*Tph2*^−/−^ and *Tph2*^+/+^ mice were born from heterozygous parents and genotyped at weaning as previously described (Alenina et al., 2009). After weaning, male littermates were kept together in a regular home-cage. At six weeks of age four mice of the same genotype were transferred to the same cage and were individually marked with unique Radio Frequency IDentification tags (RFID: 12 × 2.1 mm, 124 kHz, Sokymat, Germany, subcutaneous implantation in the scruff of the neck and under short isoflurane anesthesia). Afterwards the animals’ activity and health were regularly monitored (1h and 3h after marking and every day on the following days). At seven weeks of age, the four mice from the same home-cage were transferred for six consecutive days (120h in total) to a new version of the VBS which was designed and built for this project. Twenty-four to fourty-eight hours after leaving the semi-automated VBS, the mice were tested in the Three-chamber test (one mouse after the other) during the light phase.

### 2.3 Ethics statement

All procedures followed the national regulations in accordance with the European Union Directive 2010/63/EU. The protocols were approved by the responsible governmental authorities (Landesamt für Gesundheit und Soziales (*LaGeSo)*, Berlin, Germany). The experimental procedures were designed to allow for maximal animal welfare. Animals lived undisturbed as a group within their home-cage. Briefly, data collection was performed using automated observational methods applied to undisturbed group-housed animals. The health of the animals was monitored daily. Due to the observational nature of the study, the experimental procedure did not cause any damage, pain, or suffering to the animals.

### 2.4 Semi-automated Visible Burrow System (VBS)

#### 2.4.1 Material

The semi-automated VBS was designed and built following the description of the Visible Burrow System used in Arakawa et al., (2007), Figure 1. It consisted of a large regular home-cage (P2000, Tecniplast, Italy) into which we integrated the different compartments were built in. The cage was separated in two (open and burrow) areas by a dark wall (PVC, 35 x 9 x 0.7 cm). The open area was a square (39 x 40 x 72 cm) delimited by the dark PVC wall and three transparent walls of the cage (22 cm high) on top of which extra white walls (particle wood board with smooth white finish, 50 cm high) were added. The extra white walls kept the area bright, prevented escapes and blocked most of the outside view of the cage. On top of the open area, was set a transparent Plexiglas lid which kept the illusion of openness of the area. The lid was tilted to avoid light reflections on the videos and had ventilation holes on the side (Plexiglas 44 x 38 x 14.5 cm). In the open area, regular food chow and water were available at two apertures (12 x 4cm for food) on opposite sides (Fig. 1). The bottom of the open area was covered with bedding (0.5 cm thickness). The other area, called the “burrow area” consisted of two separated dark chambers (PVC, 8 x 13 x 6.5 cm), connected to the open surface by transparent tunnels (Plexiglas). Chamber 1 had one straight tunnel (4 x 5 x 3 cm) connected to the open area while chamber 2 had two tunnels (straight: 4 x 5 x 3 cm and L shaped: 8+13 cm long x 5 x 3 cm) leading to the open area. A black plate covered the entire burrow area (burrows and tunnels; infrared (IR) transparent acrylic glass, 18 x 38 x 0.8 cm) and the three transparent walls of this side of the large home cage were taped with black vinyl film so that the chambers and the tunnels were in near to complete darkness (Fig. 1). A grid of 24 RFID transponder readers (ID grid; Phenosys, Germany) placed under the VBS cage provided automated, continuous and simultaneous spatio-temporal information on each RFID tagged-animal present above. An infrared black and white video camera and two infrared lights were placed above the VBS cage. On the videos all animals were visible from all places in the VBS and in both light and dark phases. The ID grid and the video camera were connected to the same computer and data saved on an external hard drive for later manual analysis of the animal’s behaviours. The VBS cage could be easily disassembled/reassembled for cleaning of the parts in contact with animals.

**Figure 1.**
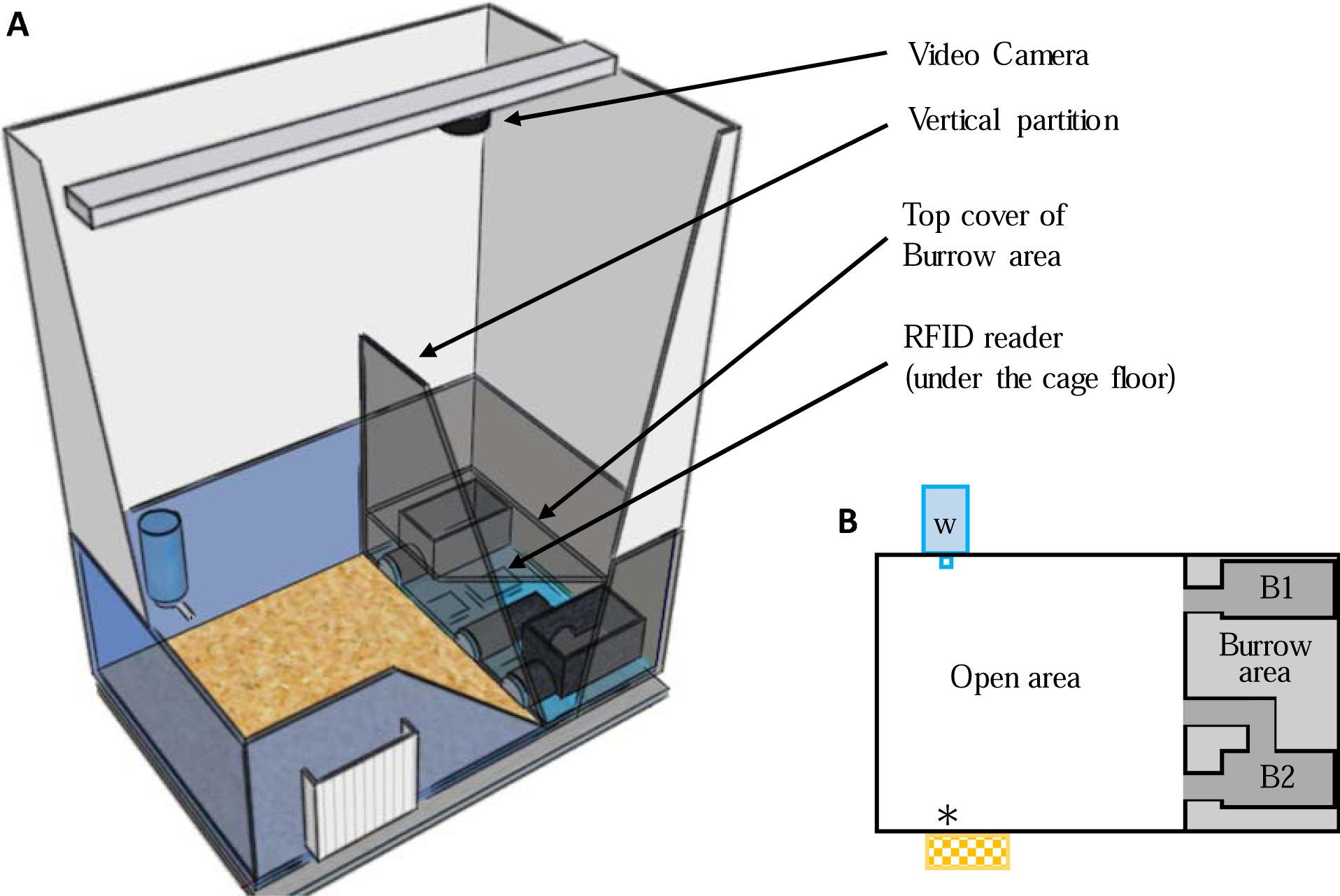
Illustrations of the semi-automated VBS. **(A)** A large rat cage was placed on top of a grid of 24 RFID readers and its walls were topped by extra high white walls. A camera was placed on top of the cage and aligned with the edge of the vertical partition (IR transparent, dark) of the burrow/open areas (drawn on SketchUp.com). **(B)** Schematic top view of the VBS cage with B1: one-tunnel burrow, B2: two-tunnel burrow, w: external water bottle with spout inside the cage, *: food access zone with outside food chow reservoir.

#### 2.4.2 Method

Each mouse was weighed before entering and after leaving the VBS cage. Each group of four mice spent five experimental days (five dark and five light phases) spanning over six calendar days in the VBS cage. They entered the VBS at the onset (or a maximum of 30 min before) of the dark phase on experimental day 1 (at 18:00) and were removed from the system at the end of the dark phase of experimental day 5 (after 06:00). In the VBS the mice were left undisturbed (e.g. no bedding change) and water and food were available *ad libitum*. The wellbeing of the animals was checked daily by inspection of the animals’ fur, posture and locomotion through the clear walls of the cage and by evaluation, on video, of their level of activity during the previous dark phase. After the VBS, mice were placed back together into the same (empty) regular home-cage.

#### 2.4.3 Data acquisition in the VBS (semi-automated)

##### 2.4.3.1 Automated collection of RFID data, videos and identification of individuals

An RFID event was automatically recorded each time the RFID transponder of a given animal was detected by a RFID reader. For each event, the control-program (PhenoSoft Control program, PhenoSys GmbH) specified the date and time, duration of the event, the identity of the detected mouse and of the activated RFID reader. Events were continuously collected and saved for the entire duration of the experiment. Thirty-second-long videos were recorded every 10 minutes during the five experimental days (CamUniversal software, Crazypixel). On each video, colored dots (one color per animal) were superimposed to the images of each mouse to allow visual identification of each individual of the group (Kolonikaefig software, PhenoSys). Marking the animals on the videos instead of color-marking their fur or ears was a more accurate, less invasive and also simpler method for long term identification of individuals within a group (Lewejohann et al., 2010, Arakawa et al., 2007).

##### 2.4.3.2 Manual annotation of behaviours (behavioural ethogram)

Similar to previous studies (Pobbe et al., 2012), only the videos of the first four hours of each phase (dark and light) of all experimental days were analyzed (25 videos per phase, two phases per experimental day and five experimental days = 250 videos to analyzed per group). Each time a mouse expressed one of the behaviours listed in Table 1, the type 1), its duration 2), where it took place in the cage 3) and the identity of any other mouse the focal mouse was interacting with 4) during this behaviour (e.g. the mouse “m1” is “chasing” for “5 sec” in the “open area” the mouse “m4”) was reported in a behavioural ethogram. One focal animal was observed at a time, all four animals were observed per video. The videos were scored by two observers (MR and AH) trained to specifically and similarly recognize the behaviours described in Table 1. The same observer scored all videos of a given group of mice. During video-scoring, the observer was blind to the genotype of the group. Consistency between observers was evaluated as follows: one observer would randomly select 10 to 20 videos of a group she did not yet annotate, score these videos and compare her results with the other observers’ results. Before all the other videos were scored, if results differed, the two observers discussed discrepancies and adjusted their scorings’ strategies accordingly.

**Table 1.**
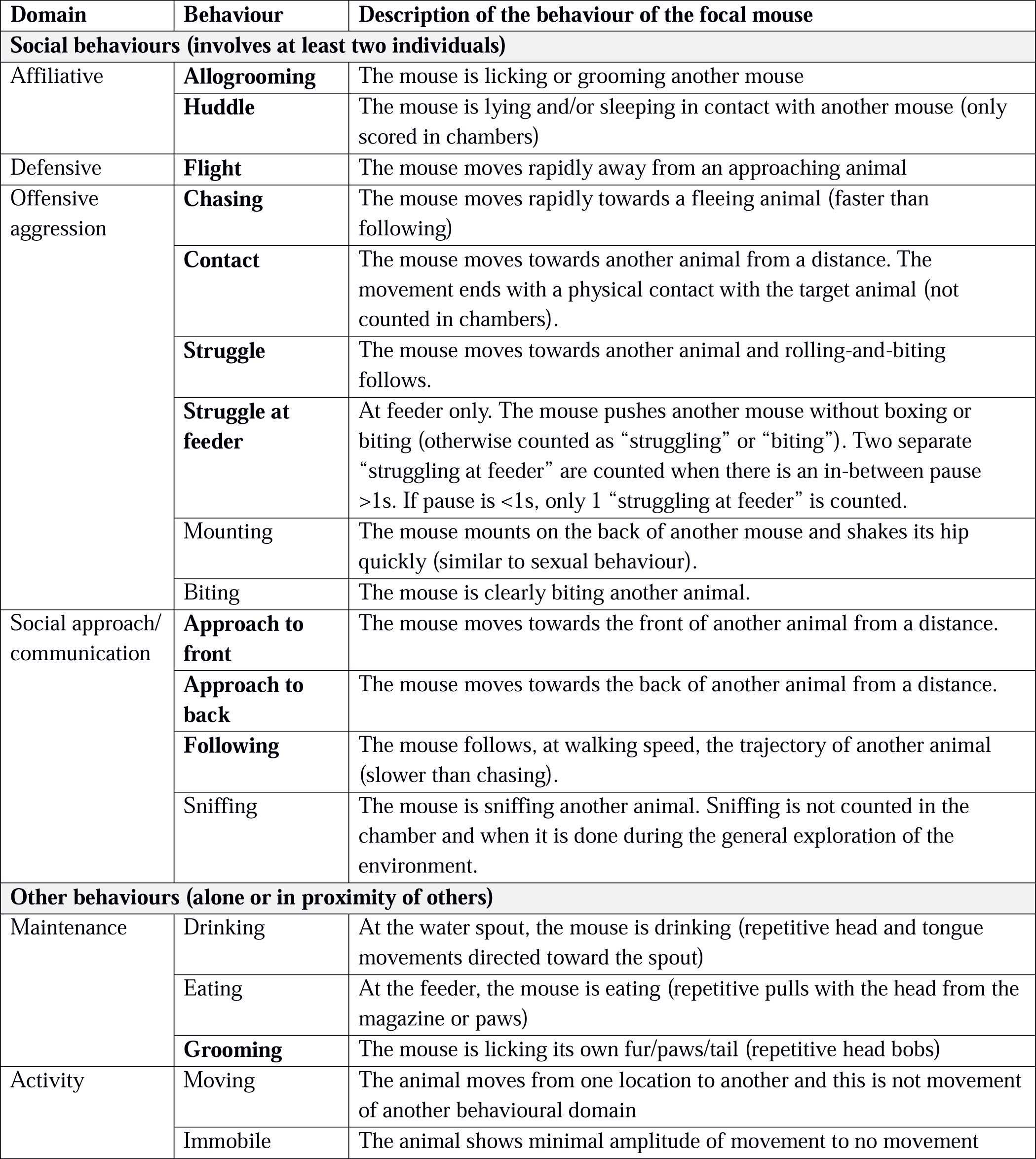
Description of social and non-social mouse behaviours. Behaviours are grouped by domains (Lewejohann et al., 2010). In bold are behaviours classically scored in VBS studies in mice (Arakawa et al., 2007), in regular font are other behaviours often scored in studies of social behaviour. The following behaviours: “being bitten”, “being sniffed”, “being mounted”, “being groomed” were scored but not analyzed to avoid redundancy with “sniffing” and “allogrooming”.

| Domain | Behaviour | Description of the behaviour of the focal mouse |
| --- | --- | --- |
| **Social behaviours (involves at least two individuals)** |  |  |
| Affiliative | **Allogrooming** | The mouse is licking or grooming another mouse |
|  | **Huddle** | The mouse is lying and/or sleeping in contact with another mouse (only scored in chambers) |
| Defensive | **Flight** | The mouse moves rapidly away from an approaching animal |
| Offensive aggression | **Chasing** | The mouse moves rapidly towards a fleeing animal (faster than following) |
|  | **Contact** | The mouse moves towards another animal from a distance. The movement ends with a physical contact with the target animal (not counted in chambers). |
|  | **Struggle** | The mouse moves towards another animal and rolling-and-biting follows. |
|  | **Struggle at feeder** | At feeder only. The mouse pushes another mouse without boxing or biting (otherwise counted as “struggling” or “biting”). Two separate “struggling at feeder” are counted when there is an in-between pause >1s. If pause is <1s, only 1 “struggling at feeder” is counted. |
|  | Mounting | The mouse mounts on the back of another mouse and shakes its hip quickly (similar to sexual behaviour). |
|  | Biting | The mouse is clearly biting another animal. |
| Social approach/communication | **Approach to front** | The mouse moves towards the front of another animal from a distance. |
|  | **Approach to back** | The mouse moves towards the back of another animal from a distance. |
|  | **Following** | The mouse follows, at walking speed, the trajectory of another animal (slower than chasing). |
|  | Sniffing | The mouse is sniffing another animal. Sniffing is not counted in the chamber and when it is done during the general exploration of the environment. |
| **Other behaviours (alone or in proximity of others)** |  |  |
| Maintenance | Drinking | At the water spout, the mouse is drinking (repetitive head and tongue movements directed toward the spout) |
|  | Eating | At the feeder, the mouse is eating (repetitive pulls with the head from the magazine or paws) |
|  | **Grooming** | The mouse is licking its own fur/paws/tail (repetitive head bobs) |
| Activity | Moving | The animal moves from one location to another and this is not movement of another behavioural domain |
|  | Immobile | The animal shows minimal amplitude of movement to no movement |

#### 2.4.4 Data analysis

An experimental day consisted of 12h of dark and 12h of light phases starting at the onset of the dark phase. An experimental day spans over two calendar days with the dark phase lasting from 18:00 of the first day to 05:59 of the next day.

##### 2.4.4.1 Activity in the VBS

The distance traveled per hour for each mouse was calculated from the event-based data generated by the ID-grid software (Phenosoft and Phenosoft Analytics, PhenoSys GmbH, Berlin). Distance traveled per hour is an indicator of the animal’s spontaneous activity over time. Due to a technical problem, data of four *Tph2*^+/+^ animals are missing from the dark phase on experimental day 4.

##### 2.4.4.2 Place preference in the VBS

Place preference in the VBS was calculated using the data generated by the ID-grid software. The relative frequency (%) of activation of each RFID-reader of the 24-RFID-reader-grid located under the floor of the VBS indicated the relative preference of the animals (averaged per phase and per genotype over all experimental days) for each of the 24 zones of the VBS. In the open area of the VBS, four zones can be distinguished: a zone with access to the feeder, a zone with access to the water spout, a “safer” zone close to the separating wall and entries to the burrow area and a more “risky”, central zone (Fig.1 and Fig. 3). In the burrow area, half of the readers were located under one burrow and its two tunnels and the other half were located under the other burrow and its one tunnel (Fig. 3). A 24-tiles heatmap represents the spatial disposition of the 24-RFID reader. The darker the color of a tile, the more the corresponding RFID-reader was activated (relative to the activation of all the other readers of the grid (%)) by the presence of animals above it and thus the greater the animals (on average) preferred this location in the VBS.

##### 2.4.4.3 Social and non-social measures in the VBS

For each behaviour described in Table 1, the total number of occurrences per animal, per genotype and per phase were analyzed (Fig.4). Potential body mass changes during the VBS housing were calculated as the difference of weight before/after VBS. Per genotype, relationships between social and non-social behaviours were evaluated using a correlation table (Supplementary Tables 1-2).

##### 2.4.4.4. Random Forest Classification for differentiation of genotypes

Machine learning was used to identify which variables, from all of the variables extracted during VBS housing (behaviours and activity, Fig. 4) were key to differentiate the *Tph2*^−/−^ from the *Tph2*^+/+^ mice. To this end we trained a Random Forest (RF) classifier (Breimann et al., 2001) using the R package “randomForest” (Liaw and Wiener 2002) and extracted the implemented feature importance (Gini index) for further analysis. We evaluated classification performance using a simple accuracy metric based on leave-one-out cross validation to ensure feasible features with regard to differentiability of genotypes. The input variables for the RF classifier were the total number of occurrences of each behaviour (illustrated in Figure 4) during the dark or light phase separately and the total distance traveled during VBS housing, for each animal of each genotype.

The estimated labels (i.e. genotype) of the test dataset were compared to the true labels of the animals and the overall accuracy of each classification, i.e. the ratio of correctly classified animals, was computed. The Gini index was automatically assigned to each variable for its contribution in differentiating the genotypes in each classification step. For robust results of the Gini index and RF we ran this procedure 100 times and reported the average (±SD) classification accuracy and Gini index. We considered behaviours with a Gini index of at least 1 as main contributors to differentiate between the genotypes.

##### 2.4.4.5. Dynamic organization of the groups

###### A. Development of aggressive and affiliative relationship strength between pairs of group living individuals using Social Network Analysis

For each *Tph2*^−/−^ and *Tph2*^+/+^ social network a node represented an animal (four nodes per network), a line between two nodes (a dyad) indicates the occurrence of at least one interaction between them and the thickness of the line (the total number of interactions between a dyad) represents the strength (weight) of this relationship (weighted directed network). The higher the number of interactions was, the thicker the edge between the respective animals. For each selected behaviour the development of its social network was evaluated during the dark phase of each day as more occurrences of behaviours happened during this phase. With such day by day network representation, we can visually illustrate dynamics and quality of interactions between pairs of individuals within their social network. The social network analysis of this study focused on the daily dynamics of overall interaction strength of both “struggle at feeder” and “allogrooming” networks in the VBS. This parameter (overall interaction strength) reflects the role of a single animal or its “relationship strength” within a network. The overall interaction strength was assessed as a node’s (a single animal) total number of interactions within the directed network (in and out). We focused the analysis on overall interaction strength instead of incorporating the directed versions of in-strength and out-strength due to the high similarity between those parameters in the observed data. For the choice of these variables and of this parameter see the Supplementary Method document.

###### B. Emergence and stability of hierarchy using the Glicko-rating method and power distribution within groups

Individuals’ social rankings established by the Glicko rating system (Glickman et al., 1999) has been found to correlates highly with other methods for dominance ranking (i.e. David’s scores and Inconsistencies and Strength of Inconsistencies (I&SI) ranking for instance in So et al., 2015). The clear advantage of the Glicko rating metric is to report on the dynamic changes in individual dominance ratings for each of the dyadic interactions within a group (So et al., 2015, Williamson et al., 2016). Briefly, the Glicko analysis calculates individual ratings based on the evaluation of the direction of the attack of each agonistic interaction between two animals. If an animal initiated a directed “struggle at feeder” behaviour, its rating increased while the rating for the losing animal decreased and all other ratings were updated accordingly to these changes in rating. The Glicko rating model (PlayerRating R-Package), is an extension of the Elo dynamic paired comparison model (Neumann et al., 2011) that did not only iteratively compute the animals’ rank but also the standard deviation of its ranking history to get an estimation of the ratings certainty, which is further used to update an animal’s ranking. Additionally, this model updated an animal’s ranking when dyadic interactions occurred between remaining animals, recognizing the group as a network being more than a sum of separated pairs of individuals. Following Williamson et al., (2016) we set the initial ranking and certainty values equal for all animals. Differing from Williamson et al., (2016) we set the ranking and certainty equal to zero to enable negative ratings to improve the visualization of development of social hierarchy. We further set the ranking update constant equal to 1, which creates little impact on the final results and still represents an accepted value for mouse agonistic interactions (So et al., 2015). The Glicko rating and power distribution analysis were performed using the data “struggle at feeder” after the social network behaviour analysis of this study.

We used the Glicko rating system to appraise 1) if similar group stratification or hierarchy was observed in *Tph2*^−/−^ groups as in *Tph2*^+/+^ groups with the final ranking of individuals spreading above and below the initial rank mark and the most dominant animals being defined as having the highest overall ranks, 2) if one distinct dominant animal could be identified at the end of the test, 3) how individual hierarchical ratings dynamically developed over time, and 4) how rapidly, in terms of the number of scored interactions, the finally dominant animal continuously received the highest rank until the end of VBS housing. Finally, we evaluated how inequitable the distribution of power could be within same-genotype groups of mice. Here the power of the dominant male was evaluated as a ratio (relative proportion) of power, defined as the difference in Glicko rating the dominant male is imposing on the first subordinate male (second highest Glicko rating score) compared to the power projected from the dominant male to the most subordinate animal (lowest Glicko rating score). A high value represents a more strongly despotic dominant male imposing relatively similar amounts of power towards all other animals. These analyses were performed on the results from the video scoring. These data were not continuously available due intermittent video recording and scoring. This may have introduced minor inaccuracies in the history of dyadic interactions, however, such effect may have been partially mitigated by the inclusion of rank certainty in the Glicko rating algorithm.

##### 2.4.4.6 Criteria for pathological aggression

For each individual, we used from the behavioural ethogram the three following quantitative parameters: (1) the latency to first attack (filtering for “struggling” or “struggling at feeder” or “chasing”), (2) the frequency and (3) the mean duration of attacks (Miczek et al., 2003, Takahashi et al., 2010). A short latency to attack associated with increased frequency and duration of attacks would suggest escalated aggression in mice (Table 2). Other qualitative aspects of abnormal aggression were more difficult to extract from our data. The body location of bites (especially to vulnerable body parts) could not be assessed in our study as very few instances of biting were observed or skin wounds were found. The lack of ritualistic behaviours (Haller et al., 2005) could only be indirectly and tentatively measured as the (4) ratio of fight/threat behaviours (Fight: struggle + struggle at feeder and Threats: chasing + following + approach to back (atb)). The theory is that at a lower fight/threat ratio, the more the threats stop escalated aggression. Any lack of responses to appeasing signals could not be evaluated from the angle (top view) of our videos. The conditions in the VBS did not allow us to evaluate if attacks were context independent, such as “aimed at the opponent regardless of its sex or state (free-living/anaesthetized/dead) or the environment (home/neutral cage)” (Natarajan et al., 2010). All the same, we could spatially locate where aggressions happened the most and if these places were appropriate places for such behaviour (Haller et al., 2005; Table 2).

**Table 2.** Pathological aggression in mice based on Takahashi et al., 2010 and Haller et al., 2005.

| | $Tph2^{+/+}$ | $Tph2^{-/-}$ | Mann-Whitney test |
| --- | --- | --- | --- |
| **Quantitative measures** |  |  |  |
| Latency to first attack, min | 153.5 ± 82.6 [158.5] | 16.1 ± 25.5 [5.2] | p<0.01 |
| Total number of attacks | 36.0 ± 24.6 [28.5] | 155.3 ± 33.6 [80.4] | p<0.01 |
| Mean duration of an attack, s | 6.8 ± 7.2 [4.2] | 6.6 ± 6.2 [3.2] | n.s. |
| Fight/threat ratio* | 6.9 ± 6.2 [5.3] | 16.2±13.6 [11.0] | p<0.025 |
| **Qualitative measures** |  |  |  |
| Where aggression occur the most | at feeder | at feeder |  |
| Bites to vulnerable body parts | n/a | n/a |  |

**Notes.** \* high ratio indicates a lack of ritualistic behaviours. Mean ± standard deviation [median], n/a: not applicable, there are no wounds at all, n.s: not significant, pvalue after Mann-Whitney test.

### 2.5. Three-chamber test

#### 2.5.1 Material

The apparatus consisted of a rectangular white box (60 x 40 x 22 cm) divided into three equal-sized chambers (20 x 40 x 22 cm). Dividing walls were made from clear Plexiglas, with rectangular openings (9 x 0.8 x 12 cm) allowing free access to each chamber. Two clear Plexiglas doors were used to block the openings when needed. Two round metal-wire grid cages with grey PVC covers at the bottom and top (Ø10 cm x 21 cm) were used. In each metal-wire cage could be placed one unfamiliar (stranger) mouse. Once the metal-wire cage was placed in the center of a lateral chamber, the opening in the wire mesh allowed the subject and the stranger mice to see, hear, smell, and touch each other but prevented fighting. A video camera above the apparatus recorded the position and behaviours shown by the subject mouse at any time and in the entire apparatus. All videos were saved on a computer for later analysis.

#### 2.5.2 Method

The subject mouse was weighed before entering the Three-chamber apparatus. The stranger mice were kept in a separate experimental room, and only transferred to the testing room when needed. The stranger mice had been previously habituated to the metal-wire cage (10 minutes daily, for at least three consecutive days before the testing day). The Three-chamber test consisted of three phases (Habituation, Social Preference and Social Recognition). In the Habituation phase, the test subject was first placed in the middle of the chamber and allowed to explore this chamber for five minutes while the access to the lateral chambers were blocked by transparent doors. Then, the lateral doors were removed, and the mouse could explore the entire apparatus for 10 more minutes. At the end of this Habituation phase, the mouse was gently pushed back into the center of the apparatus and accesses to the lateral chambers were blocked. In the Social Preference phase an empty metal-wire cage was placed in one lateral chamber and a metal-wire cage with an unfamiliar mouse that had no prior contact with the subject mouse (stranger 1) in the other lateral chamber. The location of the unfamiliar mouse in the left vs. right side chamber was systematically alternated between test animals. After the lateral doors were removed, the subject mouse could explore the entire apparatus for 10 minutes (Social Preference test). The subject mouse was then gently guided back into the center of the apparatus, accesses to the lateral chambers were blocked and the two metal-wire cages removed from the apparatus.

After an inter-test interval of 5 minutes, the lateral doors were opened, and the subject mouse was allowed to explore the entire apparatus for 10 more minutes (Social Recognition test). During the Social Recognition test an unfamiliar mouse (stranger 2) was placed in the previously empty metal-wired grid. The cage with stranger 1 was placed back into the same lateral chamber as before. The mouse had a choice between the first, already-investigated mouse (now-familiar mouse) and the novel unfamiliar mouse. At the end of the Social Recognition test, the subject mouse and the metal-wired grids were removed. The three chambers and doors were cleaned with ethanol (70%) and the metal-wired grids (emptied from the stranger mice) wiped cleaned with water and dried.

#### 2.5.3 Data acquisition and analysis

Videos of the tests were recorded and saved for offline analysis by the video-tracking system Viewer 3 (Viewer, Biobserve). In the Three-chamber test, both Social Preference and Social Recognition, were measured as the total time spent in each chamber (%) and in close proximity with the grid-cage per 5 min bins.

### 2.6. Statistical analysis

We performed two types of statistical analysis. We analysed the continuously-collected-RFID-data with regards to influence of experimental (genotype) and random (animals, batch) factors using markov chain monte carlo simulations of general linear mixed models (MCMCglmm R-package, Hadfield et al., 2010) and checked for a significant difference of the posterior distribution of the simulations with zero to assess the influence of the random variables. Other statistical tests were genotype based comparisons on a variety of experimental (distance traveled overall, per phase, per day; place preference; total behaviour occurrences; parameters of pathological aggression and weight) and analytical (social network parameters) variables. To this end we computed the exact Wilcoxon-Mann-Whitney test with multiple comparison correction. The code base is written in the R and Python programming languages and will be made available via GitHub for the published version of the paper.

## 3. Results

### 3.1. Behavioural profile of serotonin-deficient mice in semi-natural environment of their VBS home-cage across days

#### 3.2.1. Activity and Place preference in the semi-automated VBS

Animals of both genotypes showed a similar pattern of activity across days. Their activity level drastically rose and fell at the onset of each dark (high-activity) and light (low-activity) phases across experimental days, respectively (Fig. 2A). Although the activity level was higher during dark phases than during light phases, the animals were also more active during the first and the last four hours of each dark phase with a twofold decrease of activity during the four hours in between. Their activity levels were low and constant throughout light phases (Fig. 2A). Despite this similar pattern of activity, *Tph2*^−/−^ mice covered longer averaged distances over six days than *Tph2*^+/+^ animals (Fig. 2A, MCMCglmm random factors as animal and group: pMCMC=0.011, post.mean=9.865, [l-95% CI = 2.375, u-95% CI= 16.857]) during both dark and light phases (Fig. 2B, Exact Wilcoxon-Mann-Whitney test: all dark phases, Z = −2.4075, p-value = 0.01548; all light phases Z = −2.5427, p-value = 0.01031). More specifically, the *Tph2*^−/−^ mice were found consistently more active in all, except the last two, dark phases in the VBS (Fig. 2C., Exact Wilcoxon-Mann-Whitney test: day1 Z = −2.3263, p-value = 0.01954; day2 Z = −2.9485, p-value = 0.002643; day3 Z = −2.4616, p-value = 0.01319). *Tph2*^−/−^ mice were also more active than the *Tph2*^+/+^ mice on all light phases (Fig. 2C, Exact Wilcoxon-Mann-Whitney test: day1 Z = −2.2181, p-value = 0.02633, day3 Z = −2.0558, p-value = 0.04018, day5 Z = −2.1099, p-value = 0.03501). Surprisingly, after entering the VBS for the first time, the *Tph2*^−/−^ mice appeared to be less active than the *Tph2*^+/+^ mice (Fig. 2D, Exact Wilcoxon-Mann-Whitney test: first four hours of the dark phase, Z = 2.3263, p-value = 0.01954), which was not the case during the later peak of activity of the same dark phase and on the following days in the VBS.

**Figure 2.**
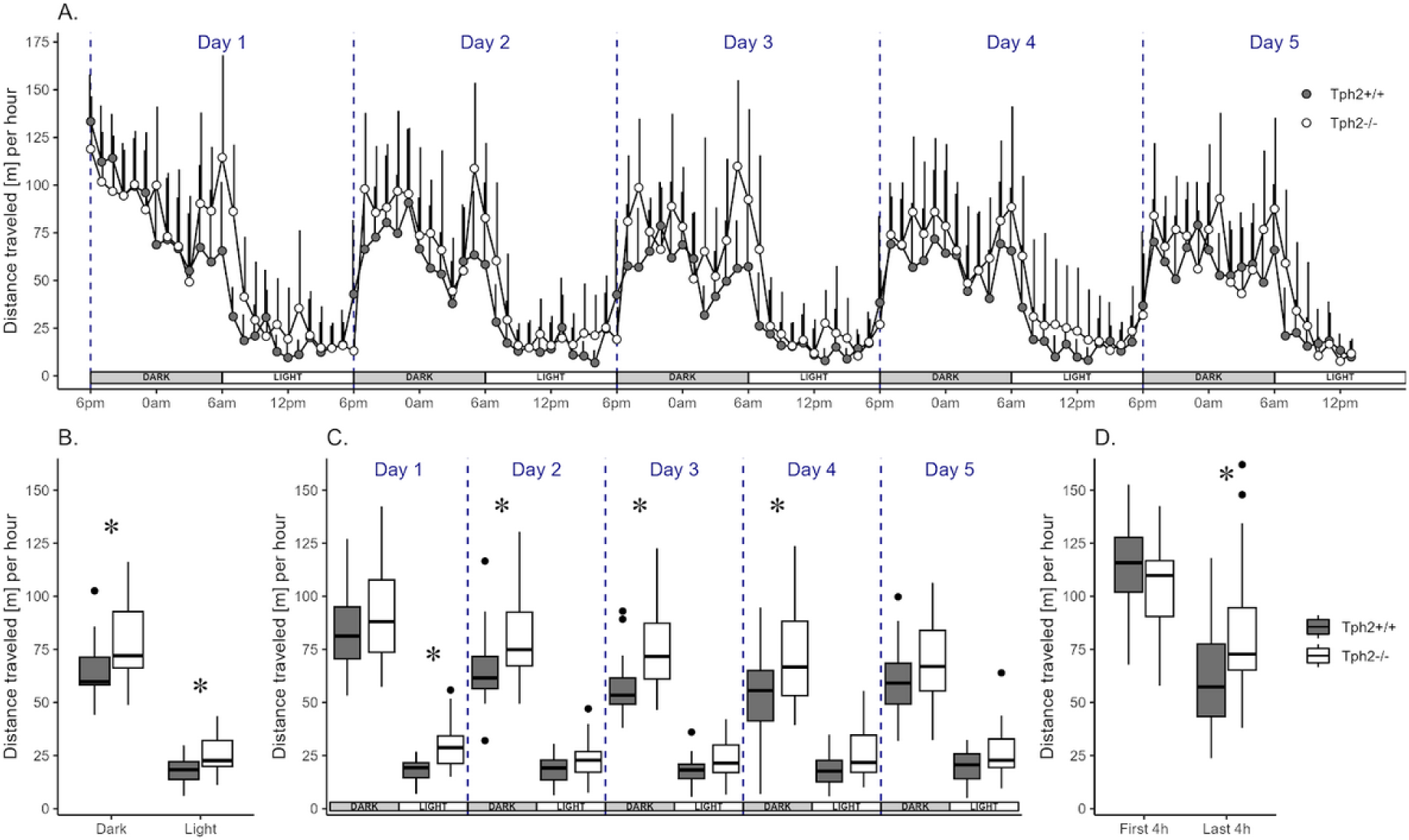
Distance traveled in the automated-VBS. **(A)** Mean distance traveled per hour (+SD) across 5 days by *Tph2*^+/+^ (solid symbols) and *Tph2*^−/−^ (open symbols) mice. Each experimental day starts (dashed line) at the onset of the dark phase (18:00-05:59; grey box) and finishes with the end of the following light phase (06:00-17:59; white box). **(B, C)** Boxplot of mean distance traveled per hour during light and dark phases, averaged over 5 days (B) or for each of five consecutive days (C). **(D)** Mean distance traveled per hour during the first and last four hours of the dark phase in experimental day 1. Boxplots show median, quartiles, 5th/95th percentiles and outlying points ({ggplot2}, R) for *Tph2^+/+^* (grey bars) and *Tph2^−/−^*(open bars). Exact Wilcoxon-Mann-Whitney test, * p < 0.05 between the genotypes. m, meter.

The heat-maps in Figure 3 revealed distinct diurnal and nocturnal spatial preferences for different zones within the VBS and between genotypes. During the light phase, animals of both genotypes were mostly detected in the burrow area with a clear preference for the two-tunnel burrow (top half of the burrow area; Fig. 3A). Although the pattern of occupation of the different zones of the VBS seemed equivalent between the genotypes (similar locations with similar shades of colors per genotype), the difference between both heat-maps (Δ = [Tph2−/−] – [Tph2+/+]) indicated that during the inactive (light) phase *Tph2*^−/−^ mice spent less time in the two-tunnel burrow and more time in the open zone than the *Tph2*^+/+^ mice (Fig. 3B, Exact Wilcoxon-Mann-Whitney test: Open area, Z = −3.8298, p-value = 8.898e-05). During the dark phase, mice of both genotypes were mostly detected at the feeder and close to the separating wall on the open side of the cage and more often in the burrow with two tunnels than in the burrow with a single tunnel (Fig. 3C). During this active (dark) phase of the day, the difference in occupation of these zones between the genotype was even more pronounced than during the light phase (stronger variations of colors on the Δ heatmap), with the *Tph2*^−/−^ mice significantly more often detected in the open area and especially at the feeder than the *Tph2*^+/+^ mice and less often in the two-tunnel burrow (Fig. 3D, Exact Wilcoxon-Mann-Whitney test: Open area, Z = −5.3018, p-value = 1.741e-10).

**Figure 3.**
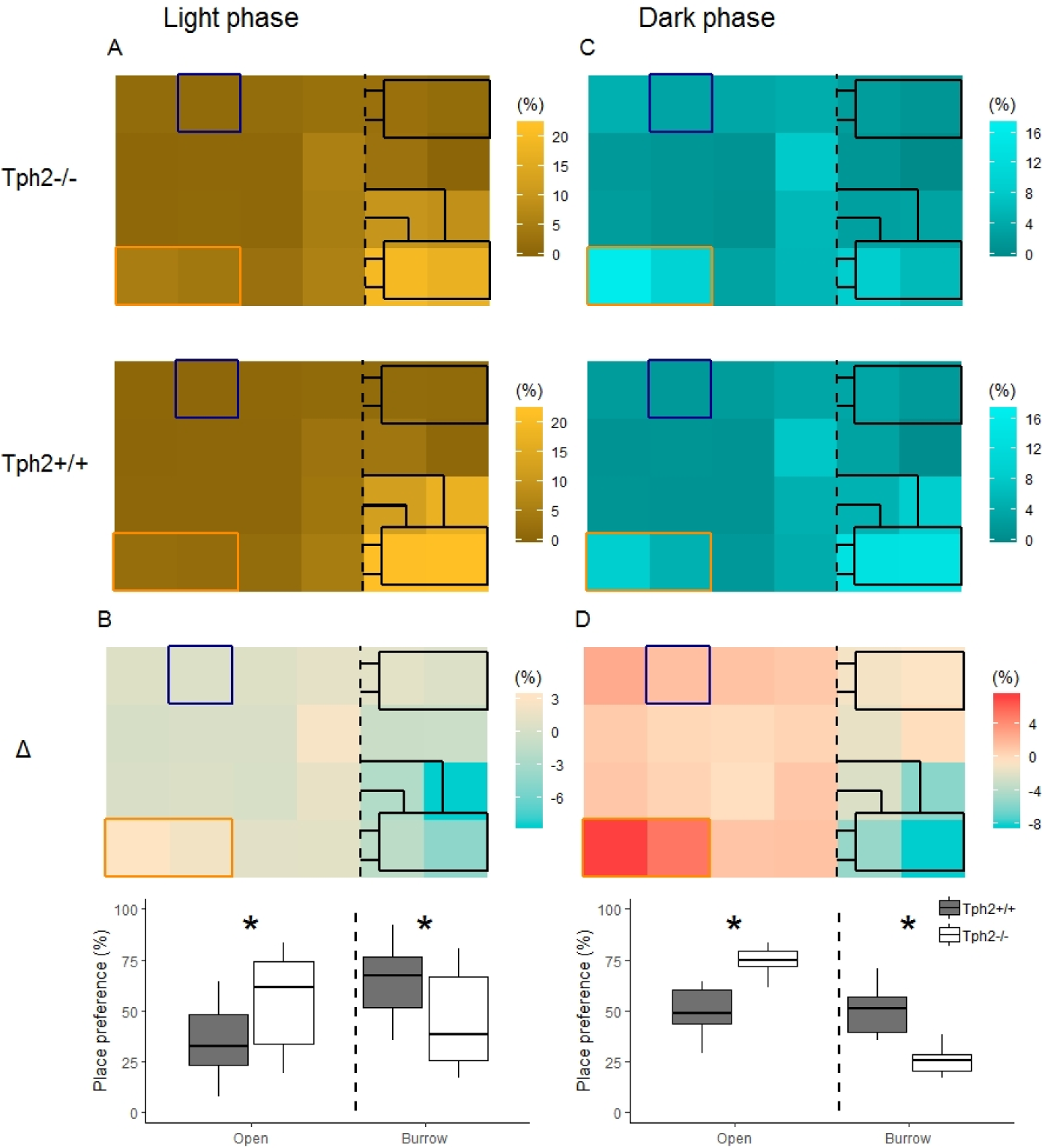
Place preference in the VBS during light (A-B) and dark (C-D) phases. **(A)** Averaged place preference (%) of *Tph2*^−/−^ (top) and *Tph2*^+/+^ (bottom) animals during the light phase (all days). Each tile of the heat-map represents the position of one RFID-reader as it was located under the VBS (24-RFID-grid). Food (orange rectangle) and water (blue square) were available from two distinct zones in the open area. The vertical black dashed-line indicates the separation between the open and the burrow areas. The brighter the color of a tile is, the more often the RFID-reader was activated (relative to the activation of all the other readers of the grid (%)) and thus the greater the animals preferred in average this location in the VBS. **(B-top)** Differences in place preference (%) between *Tph2*^−/−^ and *Tph2*^+/+^ (subtraction of heat-maps in (A)) during the light phase. **(B-bottom)** boxplot representation of averaged place preference (%) in Δ, by zone (open *vs.* burrow). **(C)** Averaged place preference (%) of Tph2-deficient (top) and Tph2-control (bottom) animals during the dark phase (all days). **(D-top)** Differences in place preference (%) between *Tph2*^−/−^ and *Tph2*^+/+^ (subtraction of heat-maps in (C)) and **(D-bottom)** boxplot representation of averaged place preference (%) in Δ, by zone (open *vs.* burrow). Boxplots show median, quartiles, 5th/95th percentiles and outlying points ({ggplot2}, R). Exact Wilcoxon-Mann-Whitney test, * p < 0.05 between the genotypes.

Despite *Tph2^−/−^*spent more time at feeder, after five days in the VBS, *Tph2^−/−^*mice gained less weight than *Tph2*^+/+^ mice (Exact Wilcoxon-Mann-Whitney test: Z = −2.6436, p-value = 0.0072; *Tph2−/−* weight: before, mean = 18.8 ± SD: 2.0g, after, mean = 18.9 ± SD: 3.4g; *Tph2*^+/+^ weight: before, mean = 19.8 ± SD: 5.1g; after, mean = 20.4 ± SD: 4.7g).

#### 3.1.2. Social and non-social behaviours in the home-cage

In the semi-automated VBS, all behaviours listed in Table 1 were seen in both genotypes. Only “biting” (occurred five times in total: three times in *Tph2*^+/+^ and two times in *Tph2*^−/−^ mice) and “mounting” (occurred six times: three times in each genotype) were very rarely seen. All behaviours, except for “huddle” which happens while sleeping, were most expressed during the active (dark) phase of the day (See light phase in Fig. S1). *Tph2*^−/−^ mice performed significantly more offensive aggression such as “approach to back”, “chasing”, “contact”, “struggle” and “struggle at feeder” than *Tph2*^+/+^ mice and in both phases (except for “approach to back” and “chasing” during the light phase, for which the occurrences of behaviours are too rare to be meaningfully quantified; Fig. 4A, Exact Wilcoxon-Mann-Whitney test: Dark phase, approach to back Z = −2.7593, p-value = 0.0048, chasing Z = −2.3208, p-value = 0.0196; contact: Z = −2.3702, p-value = 0.0169; struggle Z = −4.3708, p-value = 2.077e-06; Fig. S1, Exact Wilcoxon-Mann-Whitney test: Light phase, contact: Z = −2.0371, p-value = 0.0414; struggle Z = −4.3624, p-value = 2.657e-06). In *Tph2*^+/+^ mice, “chasing” and “approach to back” (both phases) and “struggle” and “struggle at feeder” (in light phase only) were rare behaviours (Fig. 4A, Fig. S1). *Tph2*^−/−^ mice were more defensive (“flight”) than the *Tph2*^+/+^ mice during the light phase (Supp Fig.1, Z = −2.1969, p-value = 0.0304) but not during the dark phase (Fig. 4B). Regarding social non-aggressive approaches, during the dark phase, *Tph2*^−/−^ mice exhibited significantly less “sniffing” behaviours (Z = 2.4759, p-value = 0.0124) than *Tph2*^+/+^ mice but did not differ in total number of “approach to front” or “following” behaviour during this same phase (Fig. 4C). These behaviours were rarely observed during the light phase in both genotypes and thus, were not compared statistically (Fig. S1). In both phases, *Tph2*^−/−^ mice were found eating and drinking (although “drinking” was more rarely observed, probably because of the shortness of the behaviour) significantly more often than *Tph2*^+/+^ mice (Fig. 4D: Dark phase, drinking Z = −3.1292, p-value = 0.0012; eating Z = −3.5444, p-value = 0.0002; Supp Fig.1: Light phase, drinking Z = −3.1837, p-value = 0.0013; eating Z = −4.2074, p-value = 6.191e-06) and grooming during the dark phase was less often witnessed in *Tph2*^−/−^ mice than in *Tph2*^+/+^ mice (Fig. 4D: Dark phase, Z = 2.5875, p-value = 0.0087; Supp Fig.1: Light phase n.s.). Finally considering affiliative behaviours, *Tph2*^−/−^ mice showed less “allogrooming” behaviour than *Tph2*^+/+^ mice, in both phases (Dark phase, Z = 4.3102, p-value = 3.562e-06; Light phase, Z = 2.6884, p-value = 0.0064) while “huddled” was shown as many times as in *Tph2*^+/+^ mice and during both phases (Fig. 4E, Supp Fig.1).

**Figure 4.**
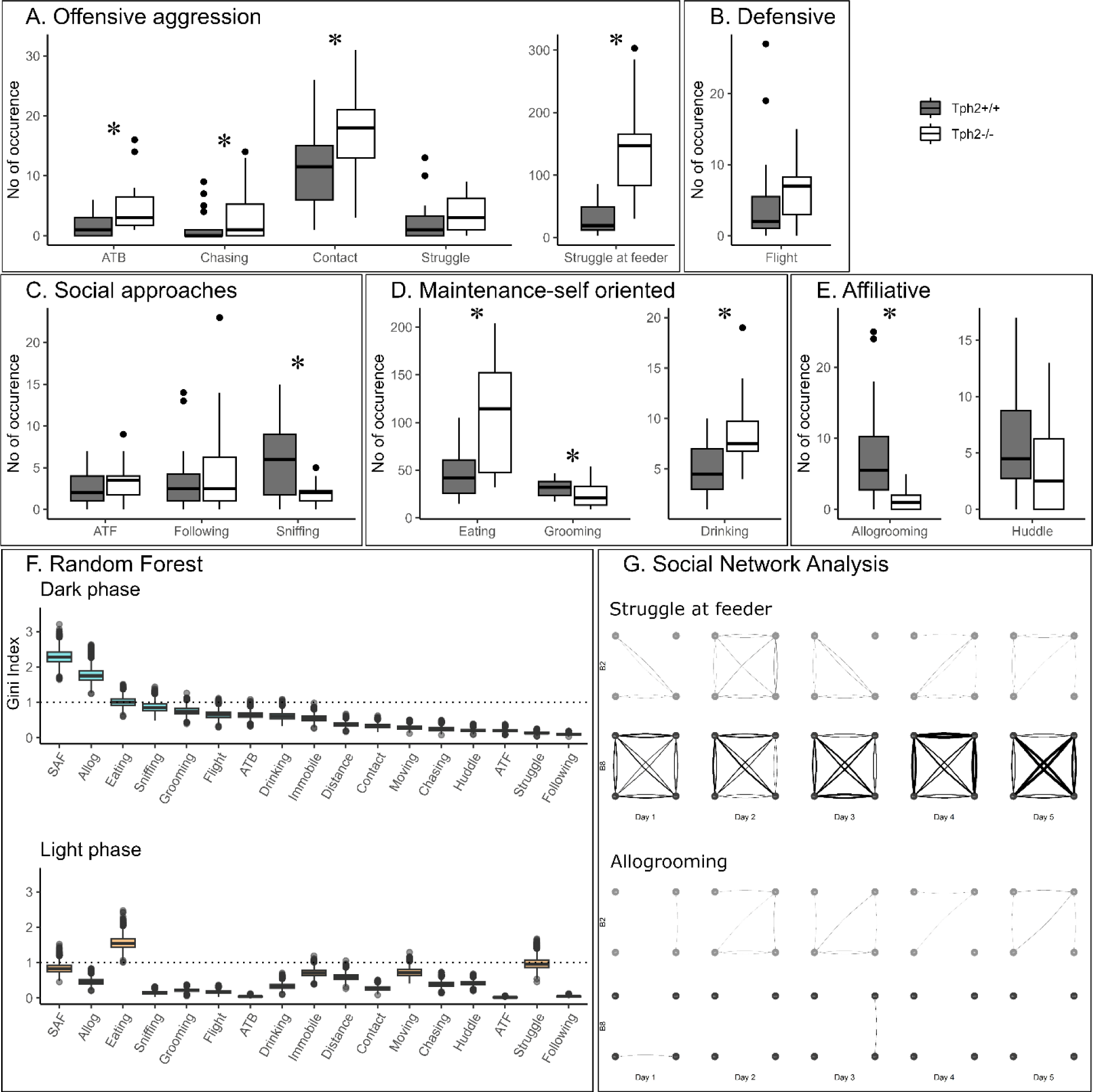
Total number of occurrences of social and non-social behaviours in the home-cage. **(A-E)** Total number of behaviours (grouped by domains) per genotype and over all experimental days during dark phases. ATB: approach to back; ATF: approach to front. **(F)** From the RF classifier, plots of one hundred Gini values, for each VBS variable, during the dark and light phases separately. Gini index >1 (dotted line) indicates behaviours highly different between genotypes. SAF: struggle at feeder. Boxplots show median, quartiles, 5th/95th percentiles and outlying points ({ggplot2}, R). Exact Wilcoxon-Mann-Whitney test, * p < 0.05. **(G)** Social networks for “struggle at feeder” (top) and “allogrooming” (bottom) of an exemplary group of *Tph2*^−/−^ and of *Tph2*^+/+^ mice across dark phases of successive days. A node represents an individual and the width of a line the overall strength of that behaviour between the pair of animals for the given day. ({igraph}, R)

The relationships between social and non-social behaviours were then explored per genotype with Spearman correlations (Tab. S1, S2). After five days in the VBS, *Tph2*^−/−^ mice did not show a weight gain different from *Tph2*^+/+^ mice that slightly increased in weight (Exact Wilcoxon-Mann-Whitney test: Z = −2.6436, p-value = 0.0072; *Tph2^−/−^* weight: before, mean = 18.8 ± SD: 2g, after, mean = 18.9 ± 3.4g; *Tph2*^+/+^ weight: before, mean = 19.8 ± 5.1g; after, mean = 20.4 ± 4.7g).

#### 3.1.3 Behavioural differentiation of genotypes using a Random Forest Classification

The training of the random forest (RF) classifier, on the social and non-social behaviours and distance traveled in the VBS of the *Tph2*^−/−^ and *Tph2*^+/+^ mice, led to a high precision in genotype prediction with an averaged accuracy of 81.4% (±1.2) over 100 runs. For each LOOCV run of the classifier we obtained the Gini index for each input variable. These values provided a robust estimate for the importance of each of the given behaviours to differentiate *Tph2*^−/−^ from *Tph2*^+/+^ mice (Fig. 4F). With the Gini index ≥1, the behaviours with the greatest potential for differentiation of the two genotypes were “allogrooming”, “struggling at feeder” and “eating” during the dark phase and “eating” and “struggle” during the light phase (Fig. 4F).

### 3.2 Role of serotonin in the dynamic organization of groups of mice in their home-cage and across days

#### 3.2.1 Evolution of aggressive and affiliative relationship strength between pairs of group-living individuals using Social Network Analysis

The network of two of the most discriminative variables between genotypes, “struggle at feeder” (offensive) and “allogrooming” (affiliative), showed clear differences in their topologies across days (Exact Wilcoxon-Mann-Whitney test, overall interaction strength, struggling at feeder, Z =-4.2158, p-value =1.2446e^−5^; allogrooming, Z=-4.6798, p-value=1.4358e^−6^; Fig. 4G showing two representative groups of mice). On a daily basis, *Tph2*^−/−^ mice compared to *Tph2*^+/+^ mice struggled at feeder with a higher interaction strength, from day 1 to day 5, (day1:Z =-3.6607, p-value =0.0001; day2:Z =-3.7961, p-value =7.3489e^−5^; day3:Z =-2.8738, p-value =0.0020; day4:Z =-5.1147, p-value =1.5709e^−7^; day5:Z =-4.277, p-value =9.4703e^−6^;Fig. 4G) but performed “allogrooming” with a lower interaction strength from day 2 to 5 (day2:Z =-5.1973, p-value =1.0111e^−7^; day3:Z =-3.3003, p-value =0.0005; day4:Z =-2.3549, p-value =0.0093; day5:Z =-3.8115, p-value =6.907e^−5^; Fig. 4G). This showed that each mouse would struggle at feeder with all the other mice of the group (between all pairs), repeatedly (edges are thick) and consistently over days, in contrast to the *Tph2*^+/+^ networks where struggling at feeder was observed between fewer and varying pairs of mice over days

#### 3.2.2. Emergence and stability of hierarchical ranking using Glicko-rating method and power distribution within groups

In both genotypes changes in individual Glicko ratings over time indicated emergence of dominance within each group (Fig 5.A for two groups and all groups in Fig. S3). In all groups, one (in two groups) or two animals’ rankings were found above their initial Glicko rating (y=0; Fig 5B) indicating in both genotypes, stratification of the individuals at the end of the VBS, into higher and lower ranked individuals (Fig 5.B). In the *Tph2*^+/+^ groups the highest ranked individual already emerged as the most dominant animal after a median number of 61 agonistic interactions (mean = 85.8 ± SD: 54.13; group scores: 16,60,61,171,121; Fig. 5C). In the *Tph2*^−/−^ groups, dominant males emerged as dominant individuals of their groups after a median number of 243 agonistic interactions (mean = 269.2 ± SD: 122.53; group scores:125,243,158,382,438; Fig. 5C). Taken together, dominance emerged more readily, after less interactions, in the *Tph2*^+/+^ groups compared to the *Tph2*^−/−^ groups. The results were consistent after controlling for the overall higher number of interactions in the *Tph2*^−/−^ groups by dividing the count at emergence by the overall count of interactions within the groups (Fig. S3).

**Figure 5.**
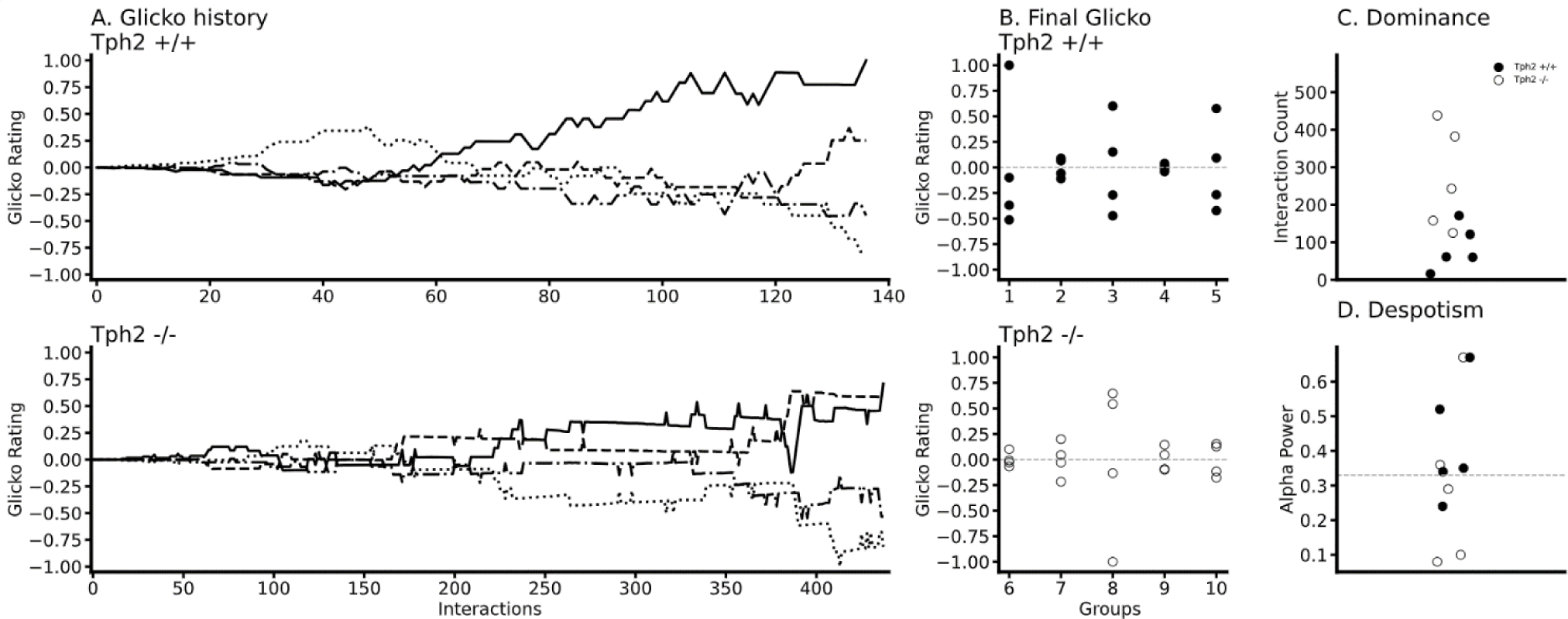
Temporal and social properties of individual Glicko ratings for *Tph2*^+/+^ groups (batches 1-5) and *Tph2*^−/−^ groups (batches 6-10). **(A)** Change in individual Glicko ratings over time of two selected groups (batch 3 (*Tph2*^+/+^) and batch 10 (*Tph2*^−/−^)) illustrating early and late establishment of dominance depending on genotype. Each line represents the ratings of one individual of the group, while the solid black line represents the final dominant male. Ratings are recalculated for every individual after each agonistic interaction. Rating values are normalized as ratio of absolute maximum rating within given group history. ({matplotlib}, Python). **(B)** Final individual ranking of each group normalized to the absolute maximum rating within a given genotype. Initial rating value is shown as vertical dotted line at zero. **(C)** Minimum number of agonistic interactions, each dominant animal engaged in before they reached the top Glicko rating that remained their rank until the end of the rating period. **(D)** Power of the dominant male defined as the ratio of absolute power exhibited from the dominant toward the second-ranked male over the total power expressed toward the lowest ranking animal. Despotic dominance is above 0.33 (dotted line).

Finally, we examined the power exerted by the dominant male on subordinate animals by evaluating how much each dominant male monopolized agonistic interactions within their social group or said otherwise, how (un)equally distributed the power was within groups. Considering the individual’s final Glicko rating, a dominant male is considered “despotic” when it imposes towards the second highest ranked male a power of one third or more of the total imposed power on the lowest ranked animal. Here we found that three out of the six most despotic males (above alpha=0.33) were *Tph2*^+/+^ animals, indicating no difference in (despotic) style between genotypes (Tab. S3).

**Figure 6.**
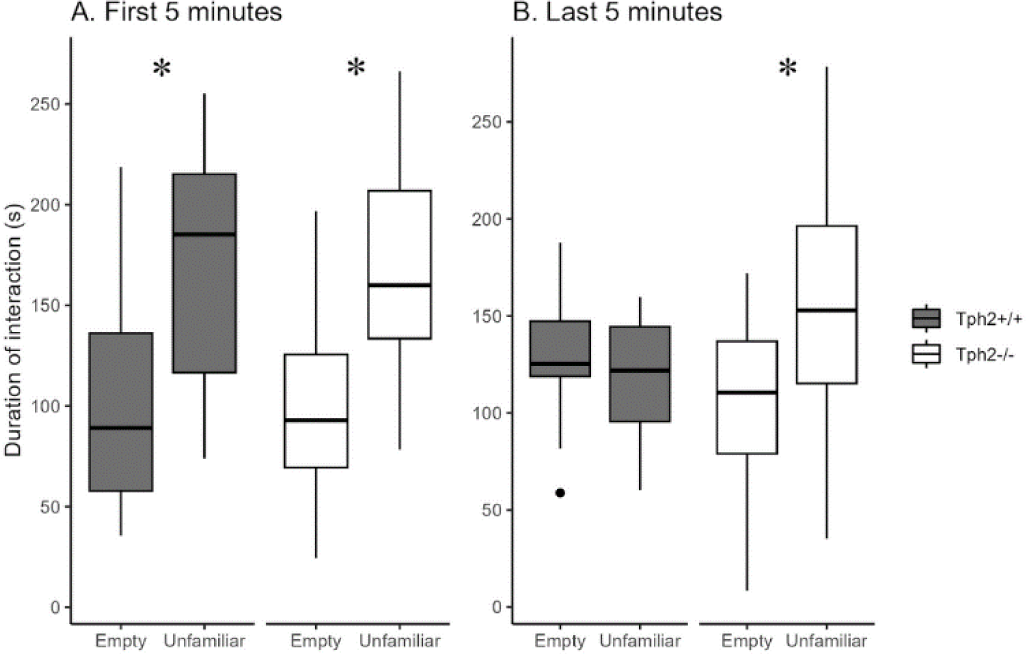
Social preference and habituation in the three-chamber test. Total duration of interaction with the empty grid or the grid with enclosed unfamiliar mouse during minutes 1-5 **(A)** and 6-10 **(B)** of test. ({ggplot2}, R). Wilcoxon-Mann-Whitney test, * p < 0.05 between empty and unfamiliar for the same genotype. s, seconds.

### 3.3 Social cognition in the three-chamber test

In the first five minutes of the social preference test both groups of mice preferred interacting with the “unfamiliar” individual (Mann-Whitney test, *Tph2*^−/−^ mice: W = 24, p-value = 0.0006612; *Tph2*^+/+^ mice: W = 16, p-value = 0.03147). In the next five minutes while *Tph2*^+/+^ mice lost interest for the now “familiar” animal, the *Tph2*^−/−^ mice keep interacting more with the mouse than with the empty cage (*Tph2*^−/−^ mice: W = 42, p-value = 0.01675). *Tph2*^−/−^ mice did not habituate as fast as the *Tph2*^+/+^ mice to a new individual. They kept investigating the novel individual over the 10 minutes of test while the preference for the novel individual was reduced in *Tph2*^+/+^ after 5 minutes of test.

### 3.4 Pathological aggression in mice

The first attack in *Tph2*^−/−^ mice occurred sooner than for the *Tph2*^+/+^ mice. They attacked (struggle and struggle at feeder together) with a higher frequency and used significantly less warning signals (e.g. threats: chasing, following, ATB) than *Tph2*^+/+^ mice. However, the duration of a given attack was not longer for the *Tph2*^−/−^ mice than *Tph2*^+/+^ mice and fights occurred mainly at the feeder (Tab. 2).

## Discussion

In this study we performed an in-depth analysis of home-cage behaviour of group living mice, to investigate the role of central serotonin in the expression of everyday-life aggression, social and non-social activities and the dynamics of group organization. In this semi-natural environment, *Tph2*^−/−^ animals did show some well conserved mouse behavioural characteristics. However, the lack of brain serotonin also resulted in significant behavioural anomalies in mice and led to altered social network characteristics and dynamics of group formation, indicative of broader cognitive impairments.

In the undisturbed ethological-like conditions of the VBS housing, *Tph2*^−/−^ mice showed typical day-night fluctuations of activity and a similar pattern of home-cage zone use throughout the day. Similar to wild type controls, they visited more frequently the food and open areas during active phases and the safe and sheltered areas during inactive phases. Serotonin ablation did not impact the typical day/night fluctuations in mouse activity observed as a response to changes in environmental light. Despite known interactions between serotonin and circadian control systems (regulation of the sleep-wake cycle) and their respective roles in the expression of seasonal mood disorders for instance (Gallardo et al., 2020), this result confirms the role of a larger neurobiological network for the regulation of these homeostatic processes. Moreover, the lack of congenital serotonin did not abolish the expression of any specific behaviour in mice, however it drastically affected their relative frequencies. Only the life-essential affiliative behaviour of “huddling” (i.e. “sleeping in direct contact with at least one other mouse”), in *Tph2*^−/−^ mice was unaffected. The expression of “huddling”, which is essential for maintenance of group cohesion (Gilbert et al., 2010, Arakawa et al., 2007), was well preserved and negatively correlated with offensive behaviours in both *Tph2*^−/−^ and *Tph2*^+/+^ groups. Finally, and despite the altered behaviour patterns of *Tph2*^−/−^ mice, their groups still organized hierarchically with time. The conservation of typical daily life characteristics along with the simultaneous expression of deficits give this model comprehensive face validity. Indeed, in their everyday life, violent patients are not socially maladapted in all contexts and all endeavors, they can also adapt to some social contexts.

Nevertheless, lack of brain serotonin had a strong impact on most other individual and group characteristics. Pathological aggression has been described as inappropriate, frequent and prolonged bouts of aggression which leads to an increased burden for the individual (Haller et al., 2005). On the five criteria of pathological aggression that we could analyze (Table 2), *Tph2*^−/−^ mice attacked sooner, more often and displayed fewer “warning” signs than *Tph2*^+/+^. However, fights were of similar duration between genotypes and rarely occurred in inappropriate zones of the cage such as in the burrows, but mostly occurred at the feeder, a typical area for fights. Moreover, *Tph2*^−/−^ mice might visit the feeder more often due to their increased metabolism (van Lingen et al., 2019), increasing their chances to meet and potentially fight as food is a resource which naturally triggers aggression (Blanchard et al., 2003). While these observations could indicate more adaptive aggression, the analysis of the day-to-day network of their fights revealed an atypical lack of de-escalation of aggression over time. In absence of de-escalation of aggression and in addition to the other markers of pathological aggression, this home cage analysis suggests *Tph2*^−/−^ mice as a potential and powerful model of pathological aggression.

On a complementary note, the lack of de-escalation of aggression in these mice could indicate poor behavioural control and cognitive flexibility. Indeed, serotonin and its multiple receptors are essential players in the control of behaviour, behavioural flexibility and the extinction of context-dependent conditioned behaviours (Bacqué-Cazenave et al., 2020, Dellu-Hagedorn et al., 2018, Alvarez et al., 2021).

Moreover, our study highlights impairments of the *Tph2*^−/−^ mice to form familiar social memories and to show typical communication behaviour. Since exhibiting adaptable and appropriate social behaviours, in particular inhibiting aggression and preventing conflict, is crucial to establish social memories and effectively communicate this knowledge to others, a lack of appropriate social knowledge could prevent *Tph2*^−/−^ mice to flexibly adjust behaviour and control aggression. While *Tph2*^−/−^ mice do not have olfactory deficits, which allows them to use olfactory cues to form social knowledge (Carlson et al., 2016, Mosienko et al., 2013), in the three-chamber test, *Tph2*^−/−^ mice kept investigating the unfamiliar mouse (three-chamber test) for twice as long as a *Tph2*^+/+^ mouse indicating a deficit in their ability to build familiar memories. The role of serotonin in forming social memories is consistent with serotonin being a new pharmacological target to counter memory alteration through lack of synaptic plasticity (Gonzalez-Burgos et al., 2008). It further plays an essential role in memory formation and especially short and working memory (Hritcu et al., 2007, Coray et al., 2022) and the encoding of familiarity and phenomena of déjà vu (Kalra et al., 2007). Moreover, and following a “for better for worse model” (Kiser et al., 2012), the lack of serotonin in the *Tph2*^−/−^ mice could dampen their sensitivity to social and non-social cues present in the environment delaying the formation of new social memories. Another key result is the lack of expression of typical communication skills. *Tph2*^−/−^ mice showed low “sniffing” and “allogrooming” counts (here and Beiss et al., 2015, Kane et al., 2012) which are also among the four most discriminating behaviours between genotypes (along with “aggression” and “feeding behaviour”) identified by the Random Forest classifier. These two behaviours are essential for social communication (Berg et al., 2018) in animals to build social knowledge (Lee et al., 2019) and for maintenance of group cohesion in mice (Wu et al., 2021, Schweinfurth et al., 2017). In mice, through their tactile sensitivity, sniffing (i.e. air movement on face and fur between animals) and allogrooming are important modalities for sharing understanding of each individual’s leadership position (Wesson et al., 2013, Lee et al., 2019). In the undisturbed environment of their home cage these animals did not display the typical behaviours that allows them to gather, communicate and use important social cues from their conspecifics. In absence of such social information, the *Tph2*^−/−^ mice could not have typical social knowledge on the different individuals of the cage which would lead to an inability for behavioural adjustment.

Here we also showed that very aggressive genetically-similar mice did dynamically organize their groups into individually stratified and stable hierarchies (dynamic and final Glicko ratings). This was although hierarchies emerged later, and the power of the alpha male was more diffuse in *Tph2*^−/−^ mice than in *Tph2*^+/+^ groups. These results highlight the non-essential role of serotonin to build up a social hierarchy but at the same time that serotonin absence impact the structuring and dynamic aspects of group formation (SNA, diffused power, late emergence of leader). The emergence of a dominant individual is a dynamic process relying in part on communication, social knowledge and behavioural flexibility of the individuals of the group (Wesson et al., 2013) which are social competences for which *Tph2*^−/−^ mice are highly impaired.

Finally, while most of the social and everyday life impairments in human patients are not easily measurable in preclinical or clinical settings, in pre-clinical research the use of ethological-like testing systems offers a novel avenue to catch everyday-life complexity. With this new semi-natural cage and the corresponding analytical tools, we developed for this project, we could study the everyday-life longitudinal symptomatology of our animal models, in different cage contexts, at both individual and group levels. This methodology provided a unique set-up for evaluating complex behaviours and further expanded our knowledge on the role of brain serotonin in pathological aggression.

## Conclusion

In this study, we show that *Tph2*^−/−^ mice present several characteristics of pathological aggression. However, beyond aggression, in their undisturbed housing conditions, *Tph2*^−/−^ mice have a more subtle, complex and dynamically maladaptive phenotype. Mice lacking serotonin had poorer communication skills (i.e. sniffing and allogrooming), possibly poorer sensitivity to environmental cues (social and non-social), altered short term memory formation of social knowledge as well as more slowly developing hierarchical ranking and different social network dynamics. With this study we highlighted the great advantages of using home cage monitoring systems for the integrated analysis of the several layers, temporality and relationships of social and non-social behaviours in mice, from individual to group levels.

We consider *Tph2*^−/−^ mice to be a great potential tool to further investigate the role of serotonin for the expression of food related aggression, short term social memory formation and aspects of social competence. Finally, with this study, by integrating home cage environments with specialized analytical methods tailored for assessing complex, spontaneously occurring home cage behaviours, we enabled the study of the dynamic and temporal aspects of mice living in self-organized groups.

## Acknowledgements

We thank Melissa Long, Susanne da Costa Goncalves, Alexej Schatz (DFG NeuroCure), Fatimunnisa Qadri, Niccolò Milani, Susann Matthes, Andrea Rodak, Lorenz Gygax, Vladislav Nachev, Milena Brunet for their support through technical assistance and knowledgeable discussion. We thank Annegret Dahlke, Monique Bergemann, Laura Rosenzweig, Bettina Müller, Reimunde Hellwig-Träger for their work with the animals. We thank Dalia Attalla, Alican Caglayan, Dow Glikman who made insightful comments on a previous version of the manuscript.

## SUPPLEMENTARY material

### Evolution of aggressive and affiliative relationship strength between pairs of group living individuals using Social Network Analysis

In this study we explored the validity of different behaviours to be represented as networks before converging on “struggle at feeder” (prominent aggressive behaviour) and “allogrooming” (affiliative behaviour) as being the most informative type of interactions for the purpose of our network analysis. Following the hypothesis of topological differences between genotypes due to increased aggression in *Tph2*^−/−^ groups, we evaluated several groupings of distinct aggression-related behaviours including but not limited to “approach to back” AND “contact” AND “chasing” but found that the reduction to single behaviours delivered more robust results. A range of standard network parameters were further explored such as clustering coefficient, betweenness centrality and shortest path length (Rubinov and Sporns 2010). The features of the current dataset however, in particular its small batch size of four animals per network, limited the information gain of including such parameters in the paper. We focused the analysis on overall interaction strength instead of incorporating the directed versions of in-strength and out-strength due to the high similarity between those parameters in the observed data.

Rubinov M, Sporns O. Complex network measures of brain connectivity: uses and interpretations. Neuroimage. 2010. doi : 10.1016/j.neuroimage.2009.10.003.

**SUPPLEMENTARY Figure 1.**
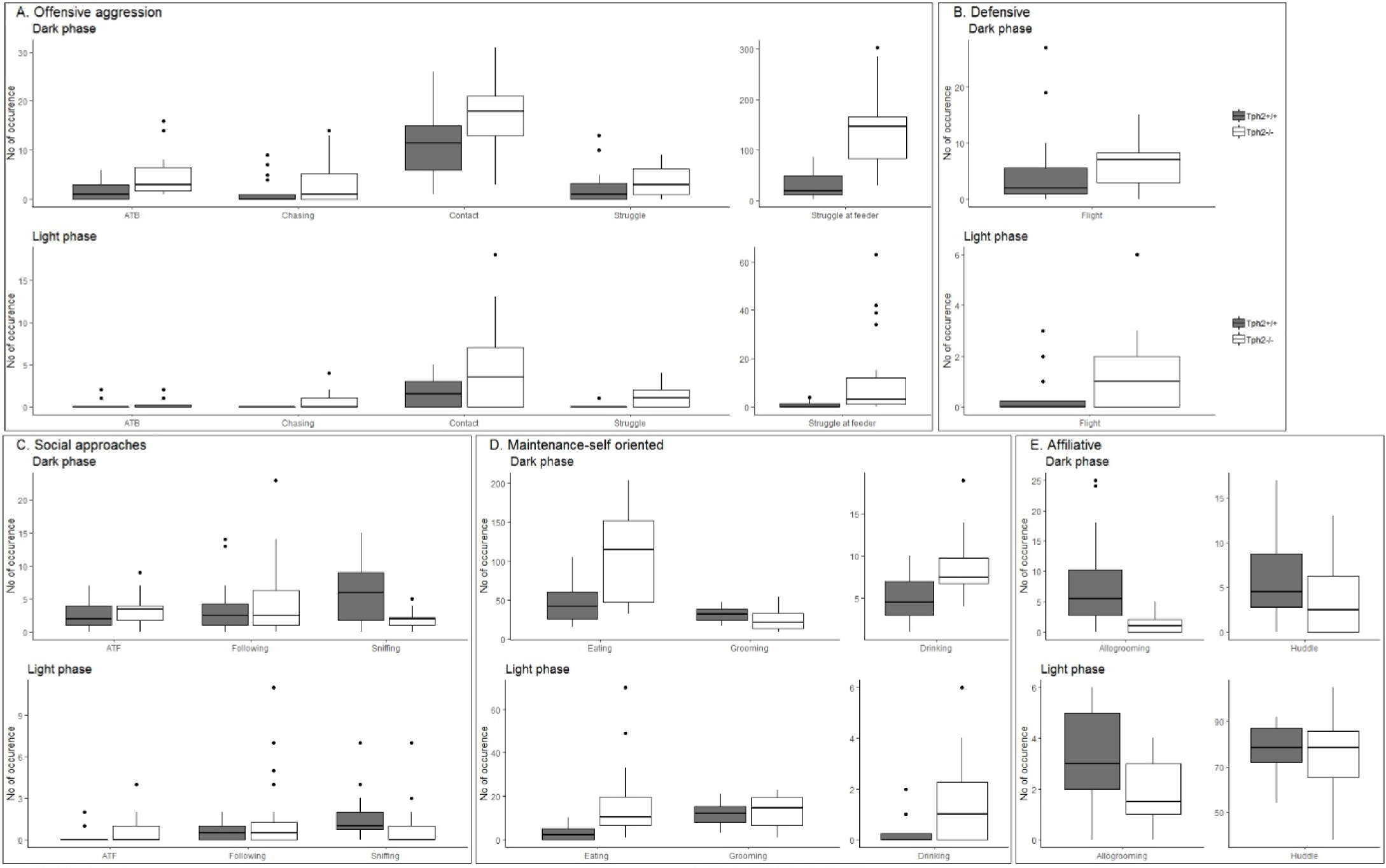
Behaviour occurrences during dark and light phases separately.

**SUPPLEMENTARY Table 1.** Correlation table of VBS behaviours of *Tph2*^+/+^ mice.

|  | chasing | contact | struggle | SAF | atb | eat | sniff | groom | drink | allogroom | huddle |
| --- | --- | --- | --- | --- | --- | --- | --- | --- | --- | --- | --- |
| chasing | — | **0.45** | **0.6** | 0.268 | 0.173 | 0.238 | 0.5 | -0.216 | 0.130 | 0.195 | **-0.676** |
| contact |  | — | **0.5** | 0.205 | **0.630** | 0.173 | 0.7 | 0.051 | 0.199 | -0.003 | -0.426 |
| struggle |  |  | — | -0.102 | 0.386 | -0.319 | 0.7 | 0.254 | -0.130 | 0.183 | **-0.713** |
| SAF |  |  |  | — | -0.012 | **0.813** | -0.2 | **-0.573** | **0.476** | **-0.653** | -0.169 |
| atb |  |  |  |  | — | -0.048 | **0.61** | -0.114 | -0.216 | 0.117 | 0.106 |
| eat |  |  |  |  |  | — | -0.28 | **-0.759** | **0.623** | **-0.629** | -0.033 |
| sniff |  |  |  |  |  |  | — | 0.156 | -0.052 | **0.451** | **-0.441** |
| groom |  |  |  |  |  |  |  | — | -0.435 | 0.415 | -0.189 |
| drink |  |  |  |  |  |  |  |  | — | -0.346 | -0.274 |
| allogroom |  |  |  |  |  |  |  |  |  | — | 0.040 |
| huddle |  |  |  |  |  |  |  |  |  |  | — |

Note. Spearman correlation; Bold <0.05 (jamovi)

**SUPPLEMENTARY Table 2.**
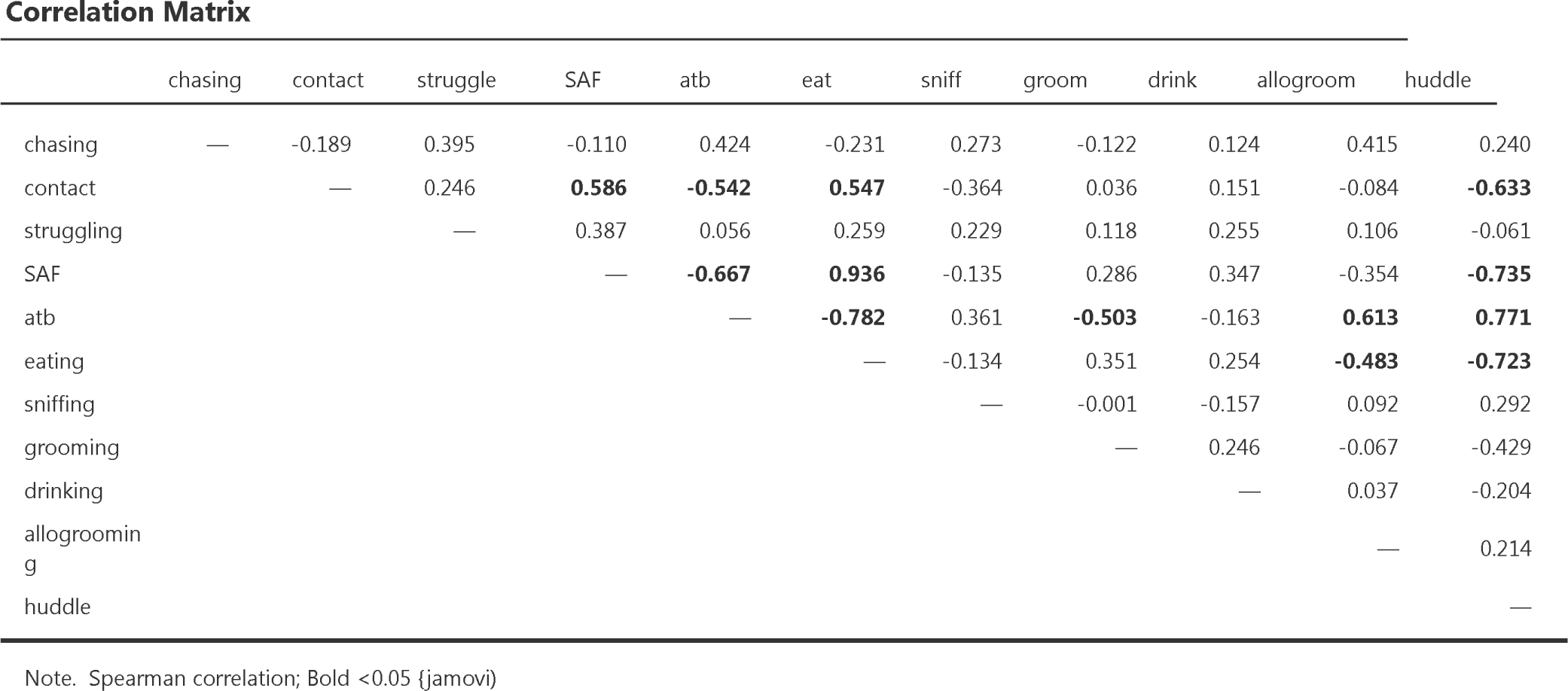
Correlation table of VBS behaviours of *Tph2*^−/−^ - mice.

**Correlation Matrix**
|  | chasing | contact | struggle | SAF | atb | eat | sniff | groom | drink | allogroom | huddle |
| --- | --- | --- | --- | --- | --- | --- | --- | --- | --- | --- | --- |
| chasing | — | -0.189 | 0.395 | -0.110 | 0.424 | -0.231 | 0.273 | -0.122 | 0.124 | 0.415 | 0.240 |
| contact |  | — | 0.246 | **0.586** | **-0.542** | **0.547** | -0.364 | 0.036 | 0.151 | -0.084 | **-0.633** |
| struggling |  |  | — | 0.387 | 0.056 | 0.259 | 0.229 | 0.118 | 0.255 | 0.106 | -0.061 |
| SAF |  |  |  | — | **-0.667** | **0.936** | -0.135 | 0.286 | 0.347 | -0.354 | **-0.735** |
| atb |  |  |  |  | — | **-0.782** | 0.361 | **-0.503** | -0.163 | **0.613** | **0.771** |
| eating |  |  |  |  |  | — | -0.134 | 0.351 | 0.254 | **-0.483** | **-0.723** |
| sniffing |  |  |  |  |  |  | — | -0.001 | -0.157 | 0.092 | 0.292 |
| grooming |  |  |  |  |  |  |  | — | 0.246 | -0.067 | -0.429 |
| drinking |  |  |  |  |  |  |  |  | — | 0.037 | -0.204 |
| allogrooming |  |  |  |  |  |  |  |  |  | — | 0.214 |
| huddle |  |  |  |  |  |  |  |  |  |  | — |

Note. Spearman correlation; Bold <0.05 (jamovi)

With some exceptions, *Tph2*^−/−^ mice presented equivalent relationships between variables than *Tph2*^+/+^ mice (correlations p>0.05). In both groups occurrence of “eating” positively correlates with the occurrence of “struggle at feeder” (Tab. S1, S2) and “struggle at feeder” (and “eating”) negatively correlated with affiliative behaviours such as “huddling” in *Tph2*^−/−^ mice and “allogrooming” and also “grooming” in *Tph2*^+/+^ group. Only in the *Tph2*^+/+^ group “huddling” negatively correlated with “chasing”, “struggle” and “sniffing”. In *Tph2*^+/+^ groups, the occurrence of a “contact” was positively associated with the occurrence of “approach to back” (ATB), “chasing” and “struggle” (Table S1, S2) while surprisingly, in *Tph2*^−/−^mice, “contact” was positively related to “struggle at feeder” (and “eating”) but negatively to “ATB” (and “huddle”) (Tab. S2). Only in the *Tph2*^+/+^ group “chasing”, “contact” and “struggle” covariated positively. In the *Tph2*^+/+^ mice “sniffing” positively correlated with “allogrooming” and “ATB” and negatively with “huddling”. In *Tph2*^−/−^ group “ATB” was the behaviour which correlated with most of the other scored behaviours. It was positively related to “huddling” and “allogrooming” but negatively to the occurrence of “struggle at feeder”, “grooming” and “eating”. Finally, in the *Tph2*^−/−^ group, “allogrooming” negatively correlated with “eating”.

**SUPPLEMENTARY Figure 3.**
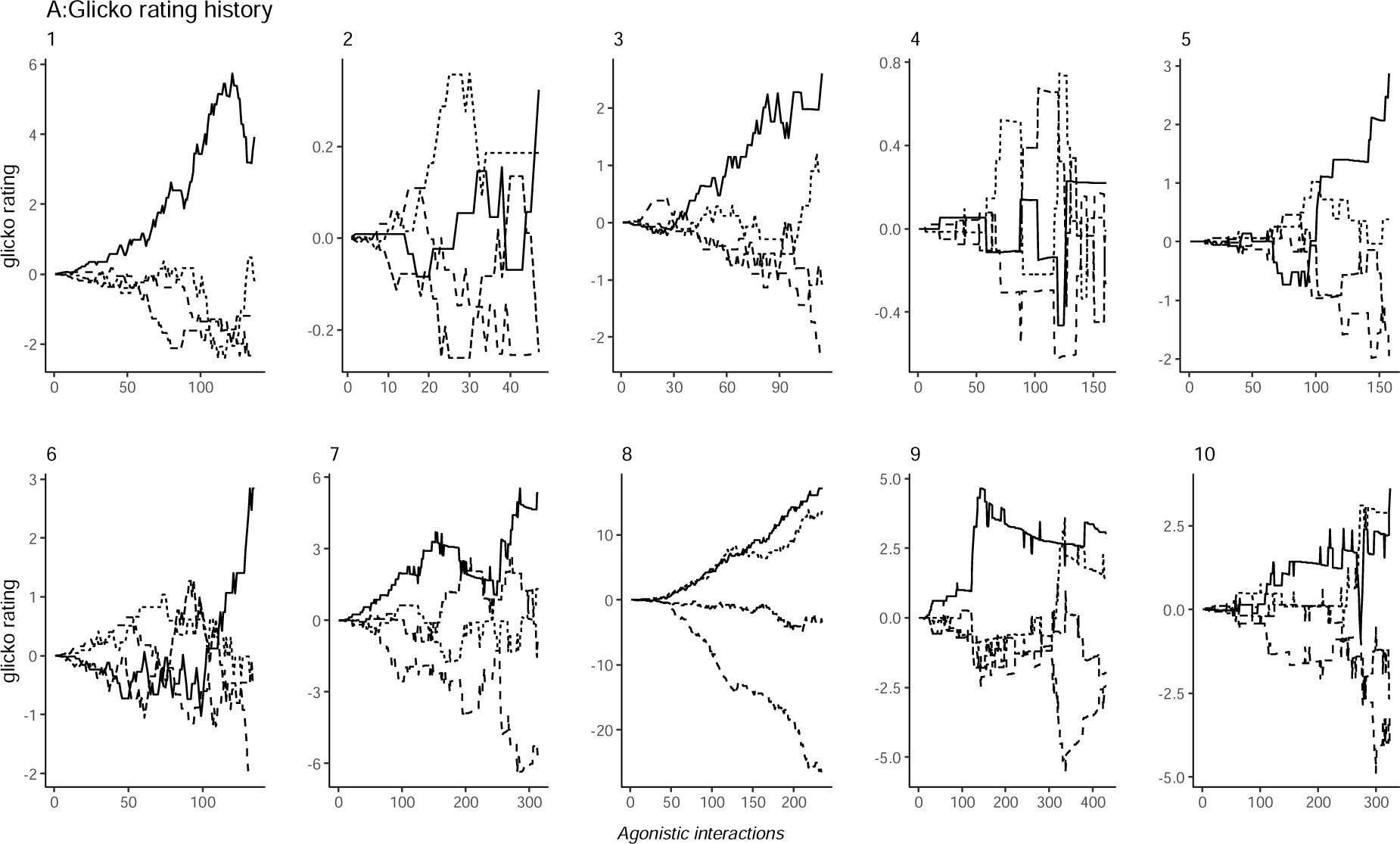
Glicko rating of all groups. Top line: *Tph2*^+/+^, Bottom line: *Tph2*^−/−^

In the *Tph2*^+/+^ groups the higher ranked individual (highest final Glicko rating at the end of a VBS stay) already emerged as the most dominant animal after a median of 68.75% of all agonistic interactions (mean: 64.46% ± SD: 34.08%; group scores: 10.19%, 100%, 44.53%, 98.84%, 68.75%). In the *Tph2*^−/−^groups, the dominant males at the end of a VBS emerged as the dominant individual of its group after a median of 80.08% of all agonistic interactions (mean: 80.3 ± SD: 12.43%; group scores: 85.04%, 73.41%, 62.95% 80.08%, 100%).

SUPPLEMENTARY Table 3.

SUPPLEMENTARY Table 3.
| **group** | **Power of dominant** | **genotype** |  |  |
| --- | --- | --- | --- | --- |
| **1** | 0.67 | CONT | Mean | 0.42 |
| **2** | 0.24 | CONT | variance | 0.02859747 |
| **3** | 0.34 | CONT |  |  |
| **4** | 0.35 | CONT |  |  |
| **5** | 0.52 | CONT |  |  |
| **7** | 0.36 | KO | Mean | 0.30 |
| **8** | 0.08 | KO | variance | 0.05766389 |
| **9** | 0.29 | KO |  |  |
| **10** | 0.10 | KO |  |  |
| **6** | 0.67 | KO |  |  |

The power of the dominant male was evaluated as a ratio (relative proportion) of power, defined as the difference in Glicko rating the dominant male is imposing on the first subordinate male (second highest Glicko rating score) compared to the power projected from the dominant male to the most subordinate animal (lowest Glicko rating score). A high value represents a more strongly despotic dominant male imposing relatively similar amounts of power towards all other animals.

